# Fixing the stimulus-as-fixed-effect fallacy in task fMRI

**DOI:** 10.1101/077131

**Authors:** Jacob Westfall, Thomas E. Nichols, Tal Yarkoni

## Abstract

Most fMRI experiments record the brain’s responses to samples of stimulus materials (e.g., faces or words). Yet the statistical modeling approaches used in fMRI research universally fail to model stimulus variability in a manner that affords population generalization--meaning that researchers’ conclusions technically apply only to the precise stimuli used in each study, and cannot be generalized to new stimuli. A direct consequence of this *stimulus-as-fixed-effect fallacy* is that the majority of published fMRI studies have likely overstated the strength of the statistical evidence they report. Here we develop a Bayesian mixed model (the random stimulus model; RSM) that addresses this problem, and apply it to a range of fMRI datasets. Results demonstrate considerable inflation (50 - 200 % in most of the studied datasets) of test statistics obtained from standard “summary statistics”-based approaches relative to the corresponding RSM models. We demonstrate how RSMs can be used to improve parameter estimates, properly control false positive rates, and test novel research hypotheses about stimulus-level variability in human brain responses.

## Introduction

Consider two potential titles of a hypothetical neuroimaging paper: (1) “Inferior frontal gyrus responds more strongly to the words‘chair,’‘house,’ and‘tree’ than to‘run,’‘pay,’ and‘speak’”; and (2) “Inferior frontal gyrus responds more strongly to nouns than to verbs”. These two titles may superficially appear to describe exactly the same set of findings, since categories like Noun and Verb are necessarily comprised entirely of individual exemplars like‘chair’ and‘speak’. Yet there can be little doubt that most neuroimaging researchers forced to choose between the two titles above would opt for the latter--which makes a far more interesting scientific statement. After all, we typically do not care about individual words like‘chair’ except insofar as they exemplify broader populations of items that share similar properties. As the psycholinguist Edmund Coleman observed over 50 years ago, “many studies of verbal behavior have little scientific point if their conclusions have to be restricted to the specific language materials that were used in the experiment” (Coleman, 1964; cf. Clark, 1973). The same is no doubt true of the stimuli used in modern neuroimaging studies.

Choosing between hypothetical titles like those above may seem purely a matter of preference--a researcher simply decides that she cares more about the underlying population than about the individual stimuli, and can then proceed to describe her results as such. But the conceptual move from stimulus-level to population-level inference is not automatically justified. It must be explicitly supported by appropriate statistical inference. In studies where a sample of participants respond to a sample of stimuli—as is the case in a vast number of fMRI studies—the correct analysis that allows generalization to both participant and stimulus populations involves fitting a mixed-effects model with crossed random effects of both participants and stimuli (Baayen, Davidson, & Bates, 2008; Judd, Westfall, & Kenny, 2012). This is not a hypothetical concern: in a review of 100 random task-based fMRI articles extracted from the Neurosynth database (see Methods for details), we found that 63/100 (95% Jeffreys interval = [53%, 72%]) used multiple stimuli in a context where generalization over stimuli was clearly indicated. Yet while virtually every fMRI study conducted over the past 15 years has modeled subject as a random factor (Penny, Holmes, & Friston, 2003), we are aware of only two published fMRI studies that have discussed the problem of unmodeled stimulus-related variance from a methodological perspective (Bedny, Aguirre, & Thompson-Schill, 2007; Donnet, Lavielle, & Poline, 2006), and know of no primary fMRI studies that have modeled participants and stimuli as crossed random factors.

The consequences of this stimulus-as-fixed-effect fallacy, as it is called in psycholinguistics (Clark, 1973), are potentially devastating. Strictly speaking, the *p*-values (or other inferential statistics) reported in the entire fMRI literature to date are valid only for the exact stimuli used in each study. The conclusions cannot be generalized to a broader population of stimuli without risking inflated Type I error (cf. Donnet et al., 2006). Previous work in other domains (and replicated here) demonstrates that this inflation can be dramatic, with the Type 1 error rate frequently exceeding 50% under realistic conditions (Judd et al., 2012; Wickens & Keppel, 1983).

Here we develop a Bayesian mixed model (the random stimulus model; RSM) that directly estimates the degree of stimulus variability in fMRI data and properly adjusts the key parameter estimates to account for uncertainty due to stimulus sampling. We then apply this model to a variety of real fMRI datasets with diverse stimulus samples and experimental designs, comparing the results from the standard statistical model that ignores stimulus variability to the results from the corresponding RSM. Our findings suggest that an unknown but possibly large fraction of published fMRI findings are likely to be false positives driven by unmodeled stimulus-level variability. We demonstrate that the magnitude of the problem can be considerably ameliorated by employing large stimulus samples and/or presenting different subjects with different stimuli -- in fact, in the limiting case where every participant receives a completely unique stimulus set, the standard model is the statistically appropriate model. Finally, we show how the stimulus-level parameter estimates produced by RSMs can be used to generate and test novel research hypotheses, opening up a powerful new method for studying the neural substrates of cognition.

## Results

### The Random Stimulus Model

Consider a hypothetical fMRI experiment in which participants view a total of 20 stimuli in a blocked design, half belonging to one stimulus category and half to another, and we are interested in the difference in neural response between these two categories (see Figure 1). In analyzing the data under the standard model, the regressors for each condition, which give the shape of the predicted neural response, are obtained by convolving an activation sequence representing when stimuli in that condition were presented with a hemodynamic response function. The standard model posits that these stimulus-level activations are all identical in amplitude for a given category (see Figure 1A)—a dubious assumption in most fMRI studies. In the RSM, we relax this assumption and allow the stimulus-level activations to have distributions of amplitudes, with the parameters of these distributions to be estimated from the data (see Figure 1B). Note that this approach is directly analogous to the way that virtually all existing fMRI packages already handle subject-level variability. That is, in addition to modeling subject as a random factor in order to account for uncertainty due to random subject sampling, we extend the same treatment to stimuli.

**Figure 1.**
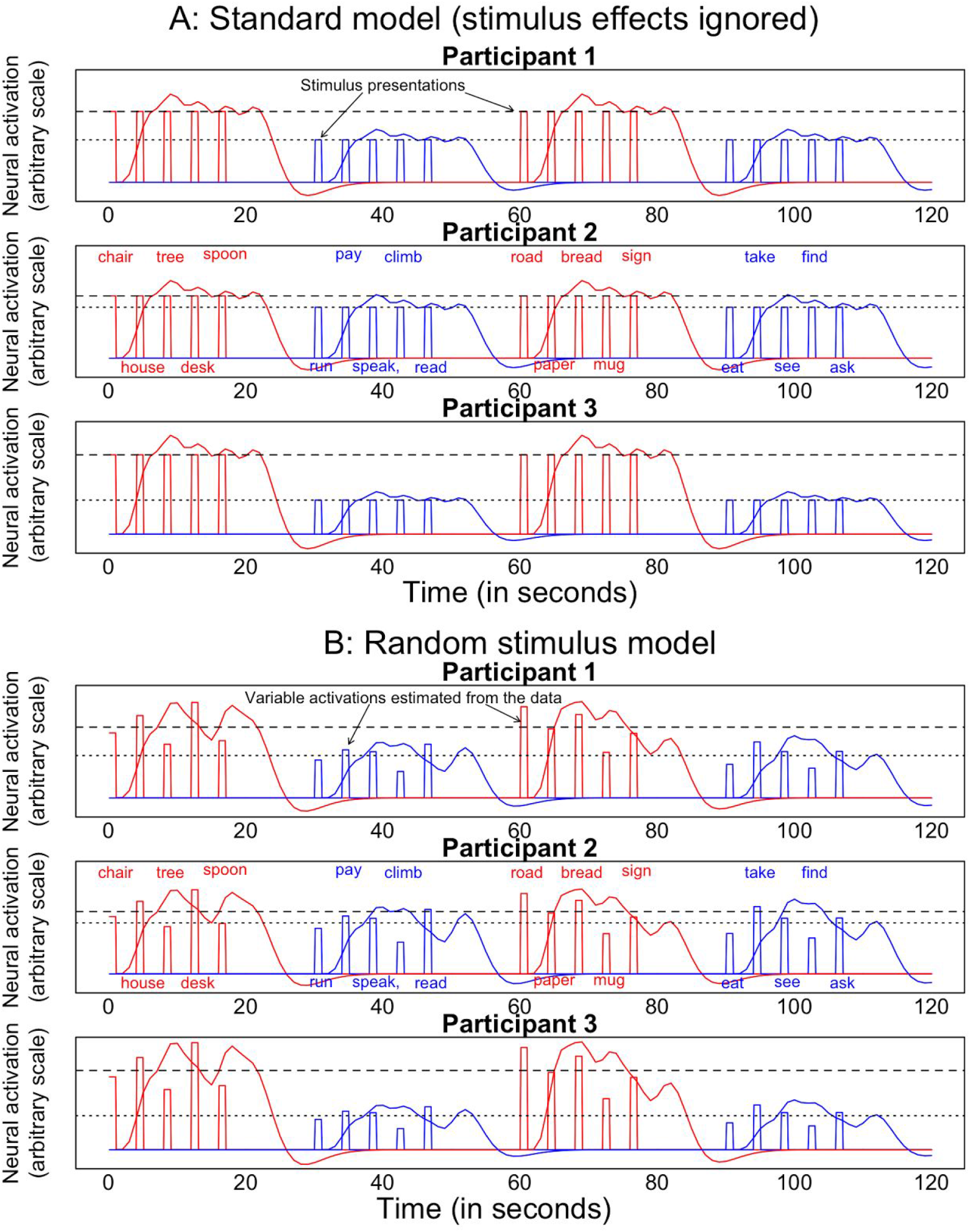
Idealized data from the standard model and the random stimulus model (RSM) for a hypothetical experiment. Stimuli belonging to one of two conditions (nouns in red, verbs in blue) are presented in alternating blocks, with the same stimuli presented in the same order to each participant. Both models incorporate subject-level variability in the magnitude of the category difference; for example, Participant 2 shows a small category difference while Participant 3 shows a large category difference. In the standard model, neural responses are assumed to be equal in magnitude for all stimuli in a category. The random stimulus model relaxes this assumption and estimates a separate response for each stimulus. For example, the noun “desk” elicits consistently high responses for all participants, while the noun “spoon” elicits consistently low responses for all participants.

To validate our Bayesian estimation approach, we conducted extensive simulations in which we systematically varied sample sizes and stimulus variabilities, fitting both the standard model and the RSM to each dataset. The full details of these simulations are given in Appendix 1. The results confirmed that the test statistics from the RSM were attenuated under the same conditions as in previously studied behavioral models (Judd et al., 2012; Westfall, Kenny, & Judd, 2014). Under the worst conditions we studied in simulation--namely, when the number of experimental stimuli is small and the degree of stimulus variance is large--the test statistic from the standard model was inflated over five-fold relative to the test statistic from the correct model. Put differently, when random stimulus effects are added to the standard model, the test statistics were reduced by over 80% in the worst case we studied. (As a preview, in some of the real datasets we reanalyze below, the reduction is even more severe than this.) Importantly, only the RSM correctly estimated the variability in the average condition difference across simulated datasets; the standard errors from both the standard model and other simpler (but incorrect) approximations consistently underestimated the true variance of the condition differences across simulated datasets.

### Does the amygdala preferentially respond to emotional faces?

To illustrate the scope and magnitude of the stimulus-as-fixed-effects fallacy in fMRI, we first focus our attention on one of the most well-established neuroimaging findings: the role of the amygdala in affective processing. The amygdala has been shown in hundreds of fMRI studies to increase activation in response to biologically salient stimuli--most notably faces--and to show a particular strong response to negative affect-provoking stimuli such as fearful or angry face expressions (Breiter et al., 1996; Morris et al., 1996). However, the number of stimuli used in studies demonstrating this effect is often small--in many cases, fewer than 10 stimuli per experimental condition. As we note above, this is precisely the situation in which inflated statistical significance is expected.

To quantify the effects of modeling stimulus as a random factor on the amygdala response to emotionally salient stimuli, we used data (n = 111 subjects) from the Human Connectome Project (Barch et al., 2013; Van Essen et al., 2013). In the HCP Emotion Processing Task--adapted from an earlier task used by Hariri and colleagues (Hariri et al., 2002)--participants view blocks of either faces (20 in total) or geometric shapes (3 in total), and make an unrelated perceptual matching judgment. The face stimuli have either fearful or angry facial expressions (10 per condition). We first analyzed the data using the standard model, where subjects (but not stimuli) were modeled as random effects. For each contrast of interest, we define a test statistic *z* = μ/σ, where μ and σ are the mean and standard deviation, respectively, of the posterior samples for the associated parameter estimate (cf. Kruschke, 2013). Consistent with previous reports on this dataset (Barch et al., 2013) and the broader literature, when analyzing the data under the standard model, we found a robust increase in amygdala activation for face stimuli relative to shape stimuli (z = 26), and a smaller but still notable increase for angry faces relative to fearful faces (z = 3.3). These results are illustrated in Figure 2 (top).

**Figure 2.**
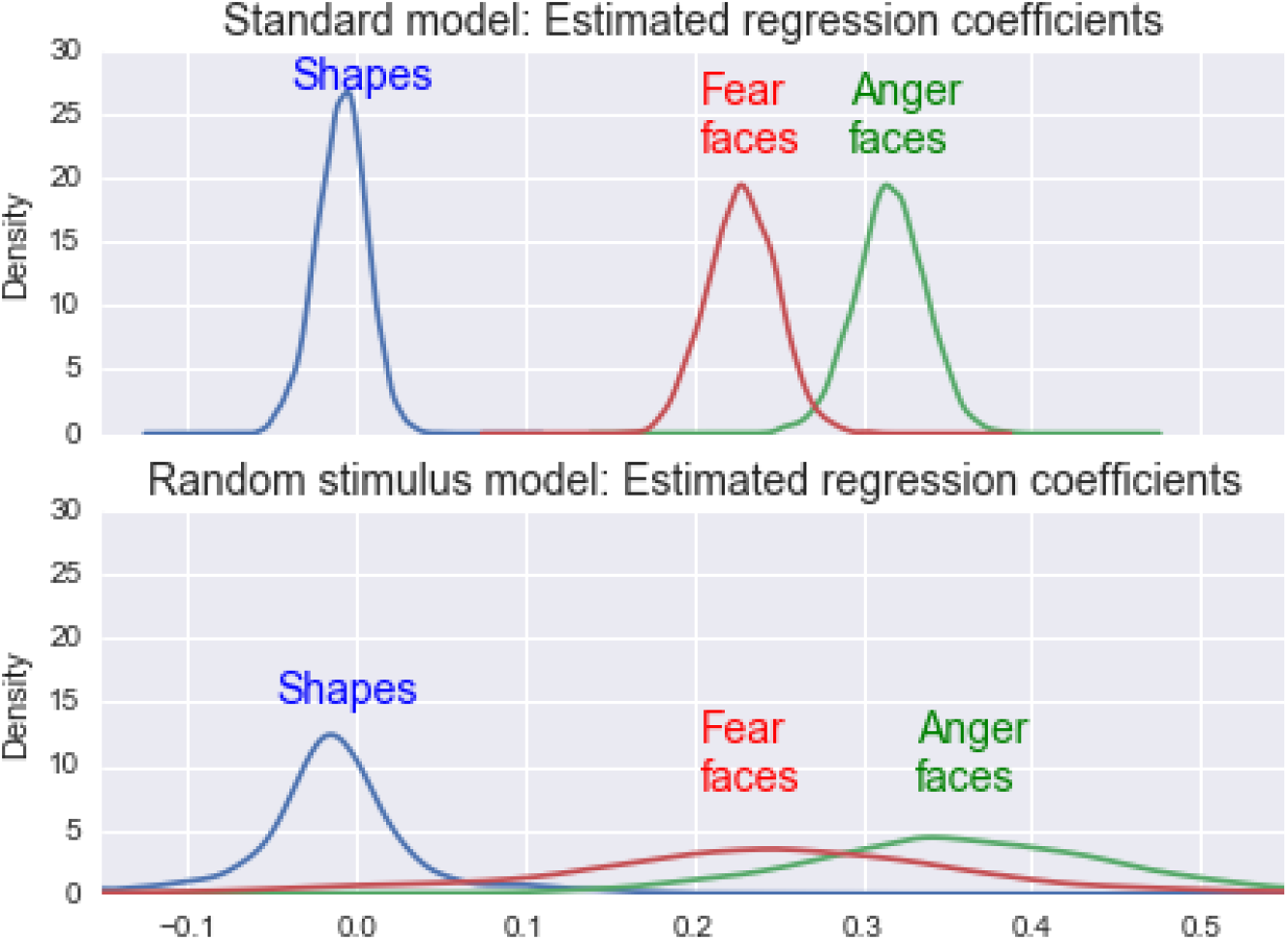
Posterior samples of the regression coefficients associated with each stimulus category, indicating the predicted magnitude of average amygdala response in response to each category, under both the standard model and the random stimulus model. Under the standard model, there is clear separation of the estimated amygdala responses toward all three stimulus categories, with anger faces evoking a somewhat stronger response than fear faces, and both face stimuli evoking a much stronger response than the simple shape stimuli. Under the random stimulus model, the means of the regression coefficients are about the same, but they are estimated with far more uncertainty. The result is that while the face vs. shape contrast is still clearly discernible, the anger vs. fear contrast is no longer distinguishable from sampling/measurement error.

As noted above, the analysis under the standard model fails to account for the uncertainty inherent in the fact that the effect of stimulus category is based on a small and highly variable stimulus sample. When we modeled the data correctly, using the RSM, we found that the test statistic for the face vs. shape contrast was reduced by 89% (from z = 26 to z = 2.8) compared to the standard model, and the test statistic for the anger vs. fear contrast was reduced by 78% (from z = 3.3 to z = 0.7). In the former case, the effect remained intact (but would have failed to do so in a smaller sample more typical of the modal fMRI study, and would not have survived multiple comparisons correction even at the current sample size). In the latter case, the remaining effect was negligible, providing essentially no basis for concluding that the amygdala responds differentially to angry versus fearful faces. These results are illustrated in Figure 2 (bottom). Thus, simply accounting for natural variability in the sampled stimuli was sufficient to turn a seemingly robust and possibly scientifically intriguing finding into an unremarkable result that probably does not merit further consideration. The striking effect of explicitly modeling stimulus as a random effect is further illustrated in Figure 3.

**Figure 3.**
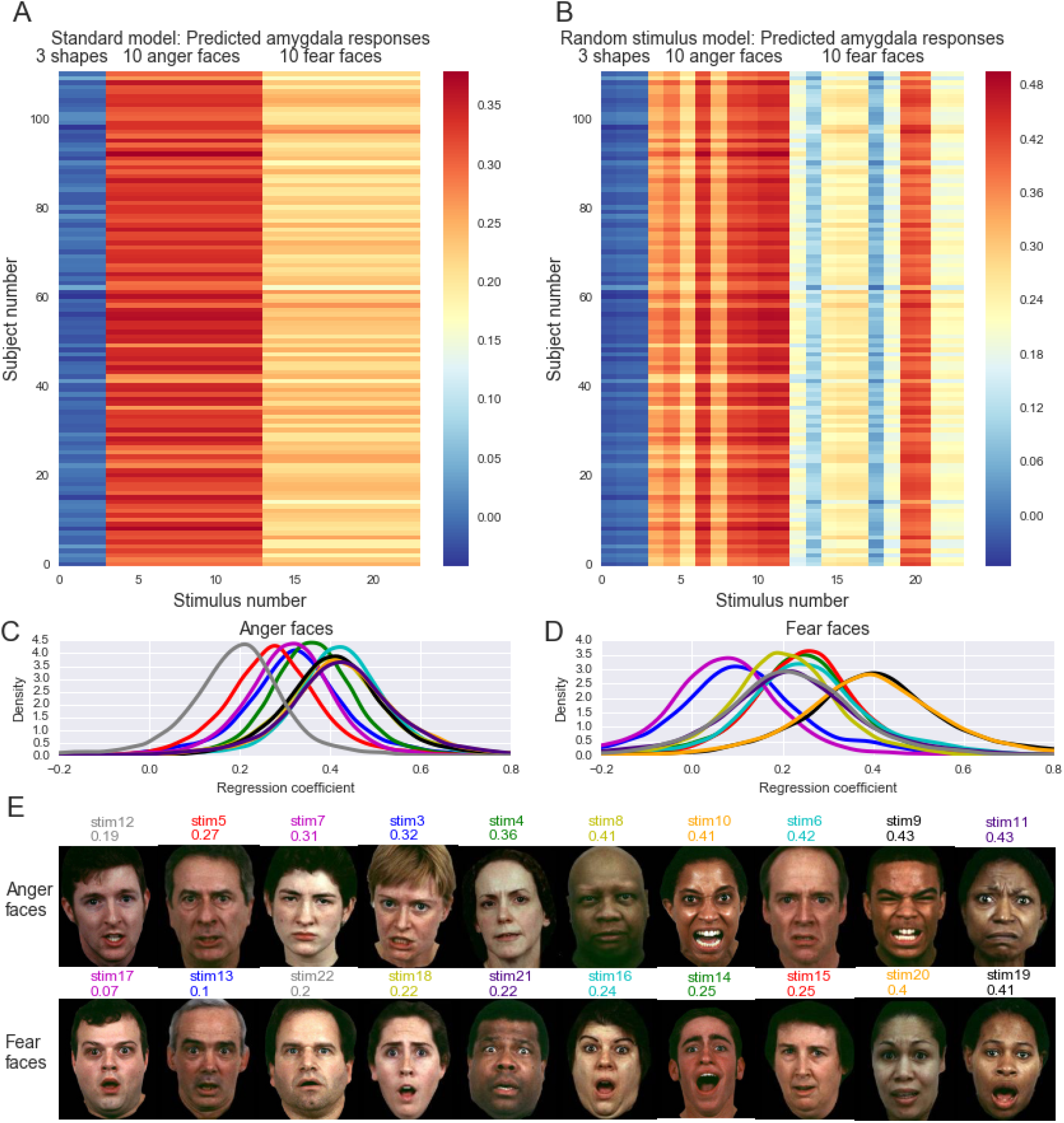
Magnitude of amygdala response predicted by the standard model (panel A) and the RSM (panel B), represented as subject × stimulus matrices, where each row (111 in total) represents a unique subject and each column (23 in total) represents a unique stimulus. Each entry of the matrix gives a (posterior mean) regression coefficient corresponding to the model’s prediction for that subject’s amygdala response toward that stimulus. Notice that the standard model assumes that a subject has the same amygdala response toward all stimuli in a given category---in other words, it assumes no random stimulus variability. While this may not be an entirely unreasonable assumption for the three relatively impoverished shape stimuli (a circle and two ovals), it is a patently absurd assumption for the faces, as a cursory visual inspection of the stimuli makes clear (panel E). When we add random stimulus effects to the standard model--resulting in the RSM--we find that random stimulus variability (evident in the within-category variance of the column means) is in fact one of the chief sources of variation in the data. The images in panel E are sorted within each emotion condition by their posterior sample means (panels C and D), which are printed below the stimulus labels.

### Whole-brain analysis reveals differential stimulus sensitivity

Next, we extended the analysis to the rest of the brain, fitting the same RSM in 100 different regions-of-interest (ROIs; we used an ROI rather than voxel-wise approach due to the computational demands of the RSM). As Figure 4 illustrates, the impact of modeling stimulus as a random factor were no less dramatic than in the amygdala for most brain regions. For the two test statistics (i.e., face vs. shape contrast and anger vs. fear contrast), the median ratios of the standard model z-statistic over the RSM z-statistic were 3.3 and 5.96, respectively, indicating reductions of 70% and 83% when random stimulus effects were added. When thresholding brain activity at even a relatively liberal threshold of z = 3, only 2 out of 100 regions (compared to 59 out of 100 in the classical analysis) remained statistically significant, and *no* region showed a significant difference between angry and fearful faces (as compared to 27 regions in the classical analysis).

**Figure 4.**
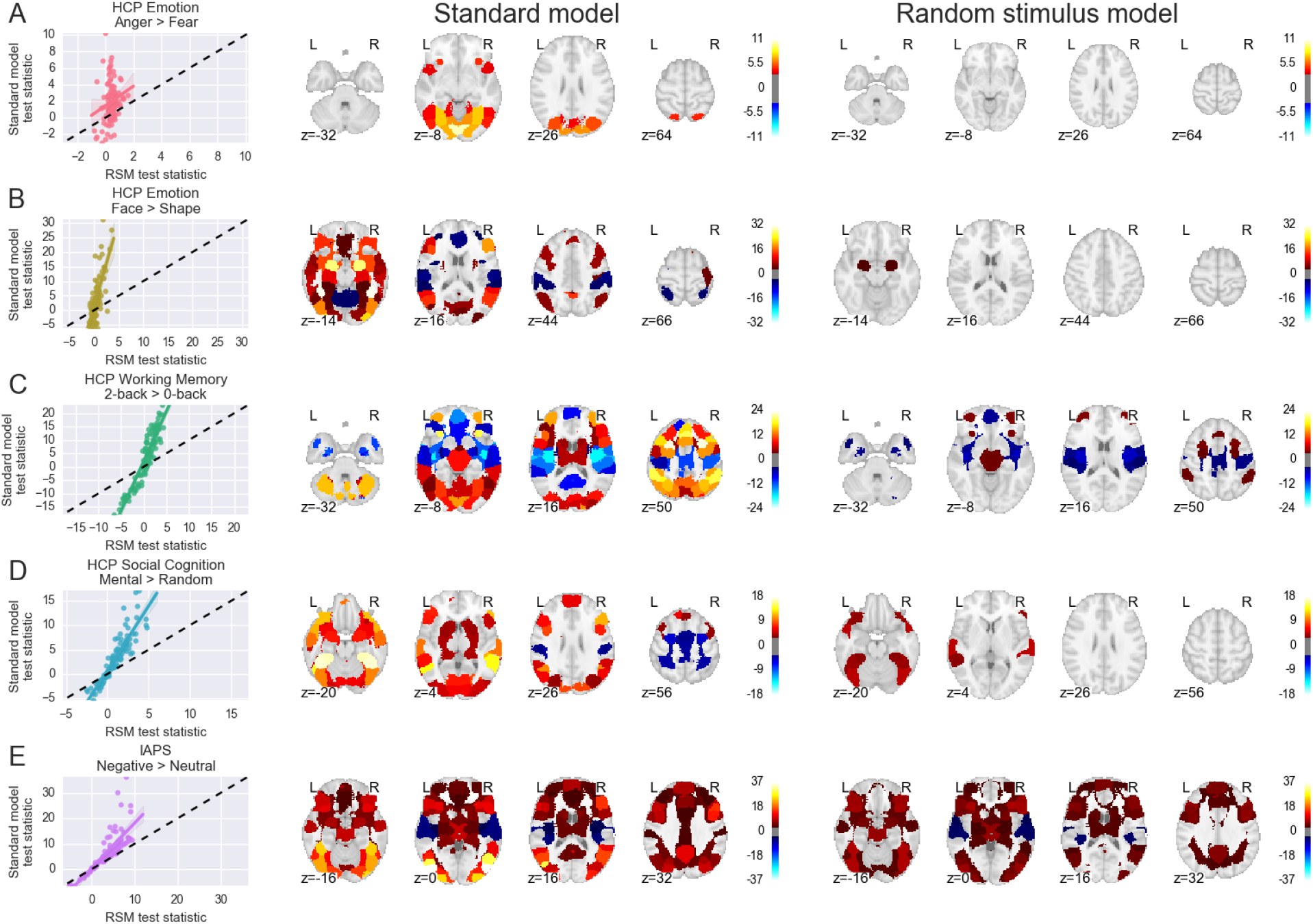
Whole-brain results for 5 contrasts from 4 different datasets when modeled with either a standard approach or a RSM. Each row displays results for a different dataset and/or contrast (see main text for details). Left column: scatter plot displaying the relationship between ROI-level test statistics from the standard model (y-axis) and RSM (x-axis). Each point represents a single ROI from a 100-region whole-brain clustering. Middle and right columns: axial slices displaying ROI-level test statistics (z statistics, defined as in the main text) from the standard and RSM analyses, respectively. Maps are thresholded at |z| > 3.3 -- comparable to using p < .001, uncorrected, in a traditional frequentist analysis -- in order to illustrate the significant drop in test statistics in most datasets when including random stimulus effects.

Intriguingly, the fusiform face area (FFA; Kanwisher, McDermott, & Chun, 1997)--which showed the most robust face-related increase in the standard model (z = 31)--failed to show a significant difference between faces and shapes in the RSM. This counterintuitive result can be understood by considering the large amount of stimulus-level variability in FFA responses to faces (Figure 5). Since the face vs. shape test statistic in the RSM depends on the ratio between the between-condition and within-condition (i.e., stimulus-level) variances, a brain region that is extremely sensitive to different stimuli of the same modality may counterintuitively fail to show a consistent difference between faces and shapes precisely *because* it is extremely sensitive (but differentially so) to individual faces.

**Figure 5.**
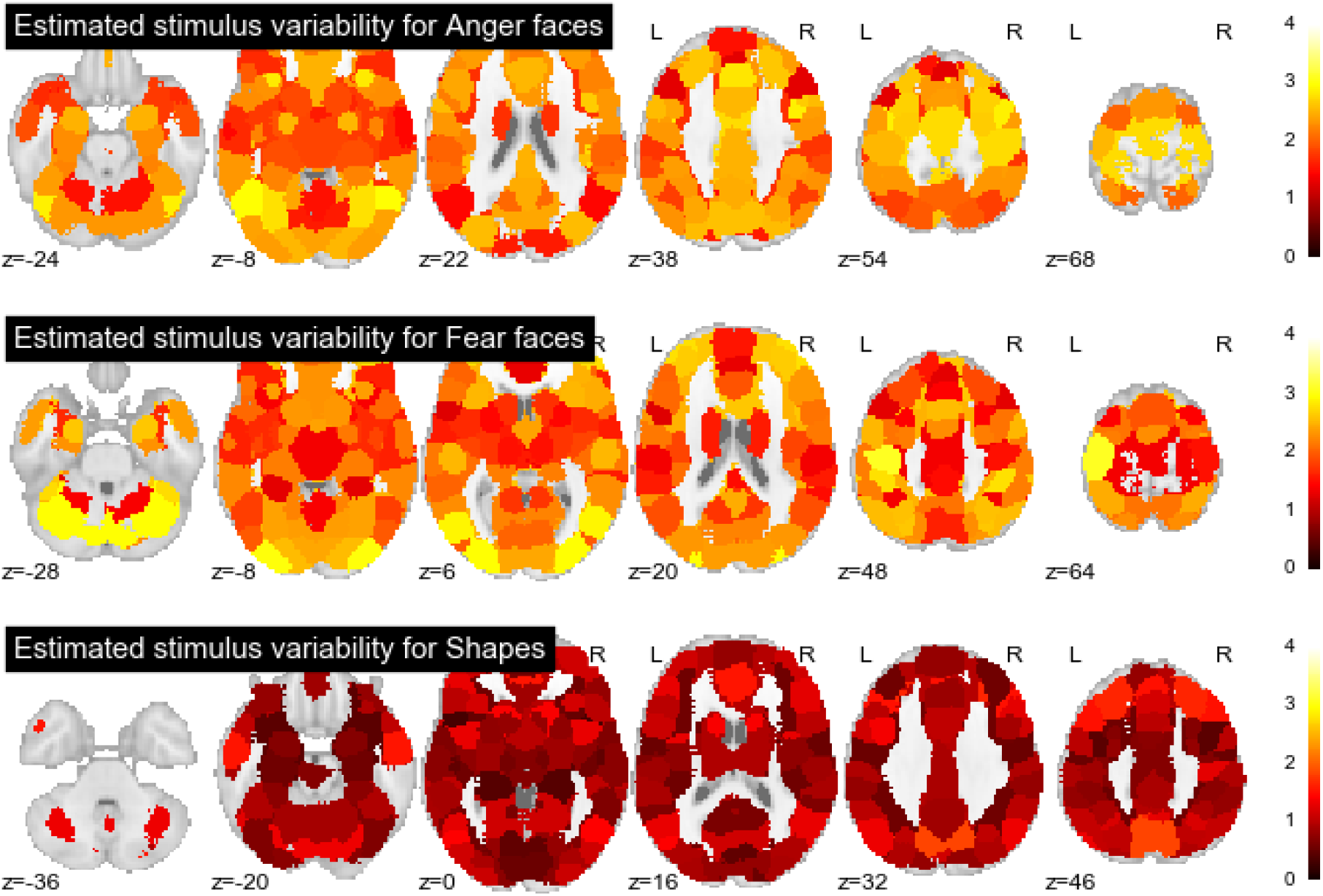
Model-estimated stimulus variability (standard deviation of the random stimulus effects) for the three stimulus categories in the HCP Emotion Processing Task, separately for each ROI. ROIs with greater estimated stimulus variability showed greater sensitivity to idiosyncratic stimulus differences within each stimulus category.

Although the resulting reduction in the test statistic may come as an unpleasant surprise, there is an important silver lining: the ability to quantify the variance in brain activity specifically related to individual stimuli provides a powerful tool for identifying brain regions sensitive to different classes of stimuli. To illustrate, Figure 5 displays the stimulus-level variability captured by the model in each brain region. Not surprisingly, stimulus-related variability was greatest in visual cortical regions; however, a number of other brain regions also showed considerable stimulus sensitivity in response to faces, including motor cortex, anterior insula (particularly for anger faces), and portions of anterior PFC. As one might intuitively expect, the variability in responses to the 3 simple geometric shapes was muted in comparison to the response to faces.

### Generalization to other datasets

To ensure that the HCP Emotion Task was not an outlier, and that our conclusions apply more generally, we repeated our random stimulus analyses on several other datasets. We fit RSM models to two other HCP functional tasks -- the Social Cognition Task and the Working Memory Task -- as well as a non-HCP emotion processing dataset (Chang, Gianaros, Manuck, Krishnan, & Wager, 2015). These tasks differed widely in experimental design, stimulus modality (video clips, images, and audio narratives), number of stimuli (10 in the social cognition task, 96 in the WM task), and putative psychological processes. Nevertheless, when contrasting the RSM with the classical model, test statistics for critical comparisons were reduced considerably in all datasets (the median reductions across all 100 ROIs ranged from 12% to 83% in the datasets we examined; see Figures 4 and 6). In general, the rank-order stability of regions was high across the two analyses in terms of the test statistics (mean *r* = .77). Thus, the global pattern of activity across the whole brain was--at least in the tested datasets--relatively conserved in the RSM, and the drop in test statistics largely reflected the increased variance of the fixed-effect estimates of the experimental conditions (cf. Fig. 2).

**Figure 6.**
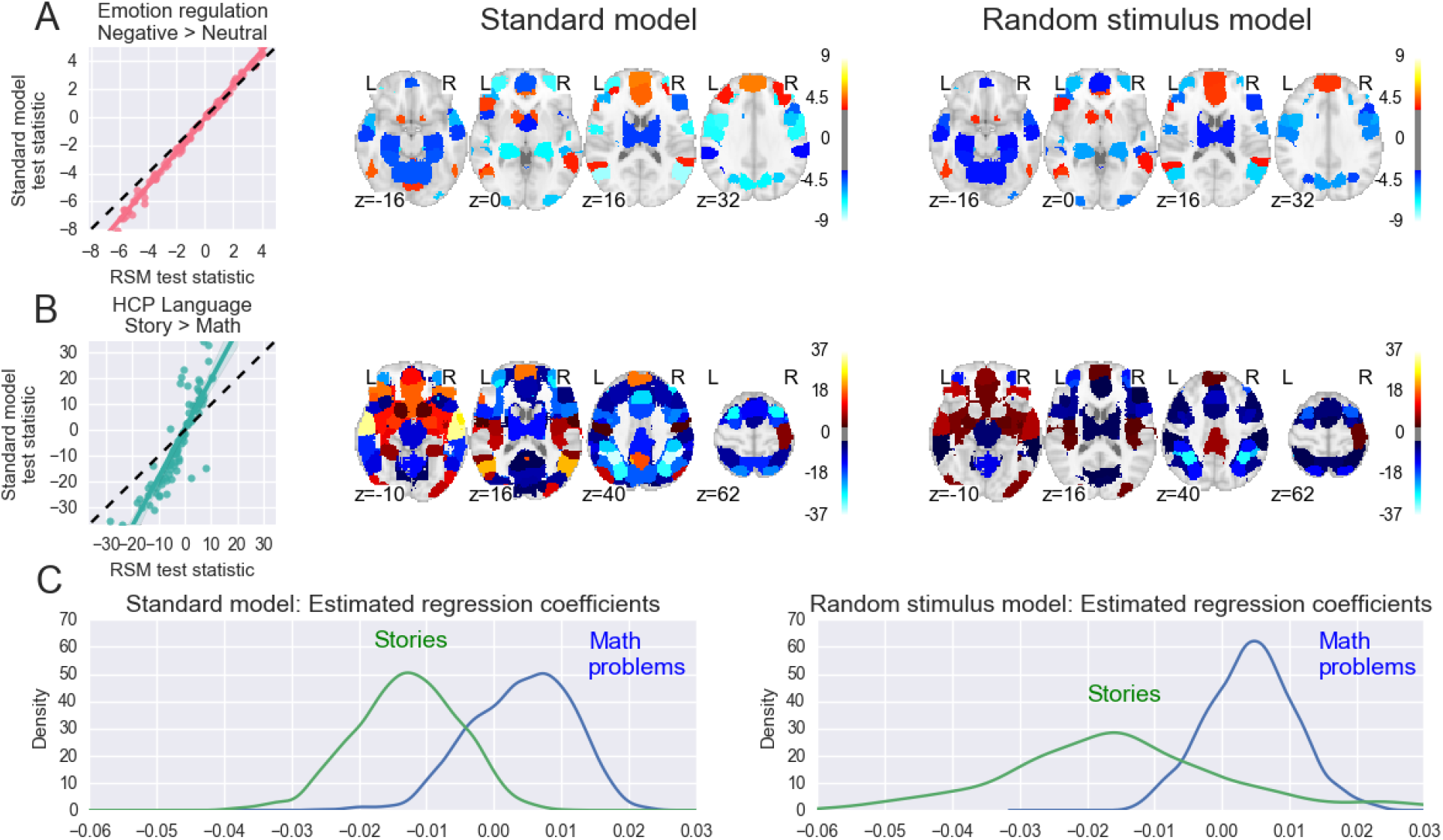
Whole-brain results for 3 datasets modeled with either a standard approach or a RSM. The interpretation is the same as in Figure 4.

### The critical role of stimulus sample size

Why were the critical test statistics from the RSM model so small compared to the standard analysis, despite the relatively large participant samples used in these analyses? The likely culprits here are the relatively small stimulus sets used in these experiments. As far as the RSM is concerned, the stimuli used in a study are just as important as the human subjects---both ultimately represent sources of random variation in the data that we would like to generalize over when estimating brain responses to different experimental conditions. Most researchers would question the wisdom of conducting an fMRI study that compared, say, 5 highly variable subjects in one condition to 5 highly variable subjects in another condition, yet many researchers routinely make essentially the same mistake when sampling stimuli. The problem is exacerbated in many cases (including in many of the present datasets) when stimuli are presented in exactly the same order to all participants---an approach that conflates order effects with stimulus effects, necessarily inflating the variance seemingly accounted for by the latter.

The important silver lining to this otherwise grim analysis is that test statistic inflation in the classical model is related in predictable ways to stimulus sample size (Judd et al., 2012; Wickens & Keppel, 1983). Thus, it should be possible to minimize the gap between the standard model and the RSM by increasing the number of stimuli in one’s experiment. To test this prediction empirically, we used two additional datasets. First, we applied the RSM to the HCP language task, which included a Math condition in which participants provided forced-choice answers to auditorily presented mental arithmetic problems. In contrast to the other HCP tasks, stimuli in the Math condition are adaptively chosen from a large set of over 7,000 candidate mental arithmetic problems based on each participant’s in-task performance. We consequently predicted that RSM estimates should be very close to standard model estimates for this experimental condition. This prediction was confirmed (Figure 6B): test statistics from the two models were very similar across the brain when comparing the Math condition to the implicit resting baseline (mean |z| = 8.47 vs. 8.12; 4% reduction). This consistency across models contrasted sharply with the large reduction observed for the Language condition (mean |z| = 4.83 vs. 2.67; 45% reduction), which presented the same 6 stimuli to nearly all subjects. As a consequence of the loss of precision in the Language condition, test statistics for the Math vs. Language contrast also showed a considerable decline (mean |z| = 13.75 vs. 5.96; 57% reduction). Figure 6C displays estimates from the standard model and RSM for a sample region (V5/MT in visual cortex), clearly illustrating the selective increase in uncertainty in the story condition.

Second, we analyzed an unpublished emotion regulation dataset (Cohen, 2009) publicly available from the OpenfMRI.org repository (Poldrack & Gorgolewski, 2015), in which 11 participants passively viewed either negative or neutral pictures. Importantly, each participant viewed 60 different stimuli (from a total set of 120). Theoretically, this “partially-crossed” design should considerably reduce the discrepancy between the RSM and classical model (Westfall et al., 2014), and this is precisely what we found: test statistics from the two models were very similar across the brain when comparing passive viewing of neutral vs. negative images (Fig. 6A; mean |z| = 2.77 vs. 2.30 for standard model vs. RSM, respectively; median ratio = 1.18). These results confirm that there is indeed a predictable and robust relationship between the stimulus sampling scheme used in an fMRI experiment and the degree of test statistic inflation one can expect to observe when using the standard (incorrect) model.

### Stimulus-level parameter estimates as a tool for exploration

While the primary reason to include random stimulus effects in one’s model is to ensure that statistical inferences can be safely generalized to new stimuli, an important secondary benefit is that this approach facilitates data exploration and hypothesis generation. The inclusion of a separate parameter for each stimulus allows one to estimate the unique pattern of whole-brain activation associated with each stimulus. Inspection of these estimates may help identify novel features of the data or design that can be subsequently tested formally.

To illustrate, consider the parameter estimates displayed for individual faces in Figure 3E. Qualitatively, there appears to be a potential trend for black faces to elicit larger amygdala responses than white faces. To formally test this hypothesis, we obtained race judgments of the 20 faces from 3 lab members blind to our hypothesis and with no knowledge of the parameter estimates. When the RSM model was recomputed with an additional fixed effect coding stimulus race, the resulting posterior estimate was suggestive of a weak race effect (z = 1.55; the other parameter estimates were all virtually unchanged). Of course, this particular analysis is circular, since the hypothesis was generated and tested using the same data. The important point, however, is that even this cursory visual inspection of the stimulus-level estimates was sufficient to suggest a scientifically interesting hypothesis that could be readily tested using independent data. Indeed, a number of previous studies (Lieberman, Hariri, Jarcho, Eisenberger, & Bookheimer, 2005) have reported race-related amygdala activation patterns consistent with our conjecture (though none included random stimulus effects, and hence the existing evidence for race effects in the amygdala is itself very likely overstated).

## Discussion

We have shown that the universal failure of fMRI studies to include random stimulus effects in statistical models can have a substantial, and almost invariably deleterious, effect on reported results. At root, the problem lies in a mismatch between researcher intent and statistical implementation: neuroimaging researchers intend for their statistical conclusions to generalize across populations of stimuli similar, but not identical to, the ones they tested; however, conventional statistical procedures only allow conclusions to be drawn with respect to the exact stimuli used in each study. Our literature survey suggests that the ramifications of this discrepancy between intent and praxis are likely to be very large: in a survey of 100 articles, we found that use of a RSM was clearly indicated in over 60% of cases. This is a conservative lower bound of the true extent of the problem, as many of the remaining studies could not have used a RSM due to otherwise avoidable limitations of their experimental design (e.g., using only a single stimulus per condition).

Given that the RSM test statistics we obtained were frequently reduced by 50% or more relative to those obtained using a standard analysis, one implication of our findings is that a large fraction of the results reported in the fMRI literature are likely to be severely inflated. Moreover, when the true mean stimulus effect is zero, variation in stimuli will generally exert some non-zero influence on brain activity (as was evident in all of the datasets we tested), which in turn will inflate Type I error. In simulations that we describe in detail in Appendix 1, we found that the Type 1 error rate for the standard model in the presence of unmodeled stimulus variability, using a nominal decision threshold of *α* = .001, ranged from about .01 to about .4, depending on the sample sizes and the level of stimulus variability. At a threshold of *α* = .05 --as would be common in many hypothesis-driven ROI-level tests--the Type 1 error rate was as high as .65, still under relatively conservative assumptions (e.g., a maximum participant sample size of 64 and a minimum stimulus sample size of 16).

Although its consequences are severe, the statistical argument that we’ve presented--based on the idea that the experimental stimuli are typically a random factor and not a fixed factor--is conceptually subtle. While the fixed vs. random distinction is not always defined consistently (Gelman & Hill, 2007, p. 245), the classical definition of a random factor from the literature on analysis of variance is that the levels of the factor that appeared in the experiment (i.e., the stimuli that were used) do not fully exhaust the the theoretical population of levels that might have been used. Importantly, this definition does *not* imply that the stimuli were selected haphazardly. Indeed, typically experimenters take great care in selecting an appropriate stimulus set to be used in the study. Rather, to say that the stimuli are “random” is simply to say that there are, in principle, other possible stimuli that could have served the experimenter’s purposes just as well as those that were in fact used. Using this definition, our literature survey suggests that a RSM is the most statistically appropriate model to use in a majority of task fMRI studies. But exceptions do exist for certain special cases. Below we list a few necessary conditions for fitting a RSM, some more conceptual and some purely statistical.

First, the stimuli in question must be inherently discrete entities, such as distinct words or photographs. An example of stimuli that, on these grounds, would *not* be modeled as random would be the varying doses of some drug administered to the subjects. While the doses are nominally discrete in that only a finite number of the range of possible doses are administered, these doses represent points on a well-defined continuum, and would simply be treated as a fixed predictor or covariate in the analysis. Second, and as mentioned above, a RSM is only indicated when the stimuli used in the study do not fully exhaust the theoretical population of stimuli that might have been used. An example of an experiment that does not satisfy this condition would be a study of brain responses toward single-digit numbers, in which the study employs all possible single-digit numbers. Third, there must be at least some degree of overlap in the stimulus sets that each subject receives. In what is overwhelmingly the most common case, every subject receives the same stimulus set, and a RSM can be estimated relatively easily. Less frequent, but not uncommon, are experiments where subjects receive different subsets of stimuli from a larger stimulus pool, but there is some overlap in the stimulus sets, such that at least some of the stimuli receive responses from more than one subject. An example would be the Math stimuli in the HCP Language task. A RSM would generally be appropriate here as well. In the least common case (6/100 of the experiments in our survey), every subject receives a completely unique stimulus set, with no overlap between sets, such that stimuli are strictly nested in participants. In this case the standard model would be a statistically appropriate model, and in fact subjects would need to respond multiple times to each stimulus for a RSM to be statistically identifiable at all.

We are aware of only two previous papers that have discussed the issue of random stimulus variability in fMRI (Bedny et al., 2007; Donnet et al., 2006). While conceptually similar, these two papers take different approaches than the one described here, and it is worth noting the differences. Donnet et al. (2006) describe a model with random stimulus effects that are different for every subject, so that the corresponding variance component is more akin to a subject-by-stimulus interaction variance component than to the stimulus variance components incorporated in our models. They also do not discuss inference on the fixed effects of activation magnitude. Bedny et al. (2007) do discuss inference on the fixed effects, but they focus primarily on conducting a separate subject-wise analysis (i.e., what we have called the standard model, which ignores stimulus variance) and stimulus-wise analysis (i.e., the conceptual complement of the subject-wise analysis, which includes stimulus variance but ignores participant variance). This approach is common in psycholinguistics, and it is certainly a step up from running only the standard or subject-wise model, but it is not equivalent to the full, correct model with crossed random effects of participants and stimuli. In particular, it does not succeed in maintaining the nominal Type 1 error rate (Raaijmakers, 2003; Raaijmakers, Schrijnemakers, & Gremmen, 1999).

What can researchers do to address the stimulus-as-fixed-effect fallacy? Broadly speaking, there are two possible strategies. The best practice approach to address the issue is to explicitly include random effects in one’s model for every factor a researcher intends to generalize over. In the present study, we used MCMC sampling to fit our mixed-effects models; however, other approaches (based on maximum likelihood estimation, variational inference, etc.) are also available (Bates, Douglas, Martin, Ben, & Steve, 2015). The primary downside of an estimation-based solution is that it is computationally intensive and may be technically demanding. At present, no major fMRI analysis package supports RSMs of the kind we employ here, limiting the ability of most researchers to produce correct inferences. While we have open-sourced the *nipymc* Python package we used to fit the models reported here (http://github.com/PsychoinformaticsLab/nipymc). this should be viewed as a provisional (and not particularly scalable) solution until more robust and widely-used packages such as FSL, SPM, or AFNI introduce support for random stimulus effects.

Alternatively, a less effective but much simpler approach to the problem is to use as large a stimulus sample as is practically feasible. In previous work, we have shown that the number of stimuli can impose a hard cap on statistical power in the RSM: when stimulus samples are very small, it may be impossible to obtain statistically significant estimates of the fixed effects, no matter how many thousands of subjects one samples (Westfall et al., 2014). This is evident in the present findings, where test statistics from fMRI experiments with only a few stimuli (such as the HCP Emotion and Social Cognition tasks) showed precipitous drops in the RSM, whereas those generated by designs with many stimuli are often negligible (e.g., Figure 6). The primary (and significant) downside of a stimulus-maximizing heuristic is that it is only a heuristic--there is no guarantee that the resulting test statistic will closely approximate the one that would have been obtained through the explicit inclusion of random stimulus effects. In particular, if the degree of random stimulus variability is large, then a huge number of stimuli may be required before the two sets of test statistics closely converge. Nevertheless, in the absence of analysis tools capable of correctly modeling multi-stimulus designs, we strongly encourage researchers to always include as many stimuli as possible in their designs. Importantly, unlike increases in participant sample size, adding stimuli rarely incurs any additional cost. Researchers can usually easily increase the number of stimuli by either (a) eschewing repeated presentation of a few stimuli in favor of single presentation of many different stimuli, or (b) using a “partially crossed” design where each participant responds to a different subset of stimuli (Westfall et al., 2014). These approaches allow one to enjoy the statistical power benefits of a large stimulus sample without increasing data collection requirements.

## Methods

### fMRI Datasets

All analyses used publicly available data obtained from one of three sources. The HCP analyses used task fMRI data from the Human Connectome Project’s “100 unrelated subject” release, accessible via the online Connectome Workbench (http://www.humanconnectome.org). These data were provided [in part] by the Human Connectome Project, WU-Minn Consortium (Principal Investigators: David Van Essen and Kamil Ugurbil; 1U54MH091657) funded by the 16 NIH Institutes and Centers that support the NIH Blueprint for Neuroscience Research; and by the McDonnell Center for Systems Neuroscience at Washington University. All HCP analyses used the preprocessed data release. No further processing of the time series was performed prior to region-based averaging and mixed-effects modeling (see below). Methods have been previously described in detail in (Glasser et al., 2013).

The emotion regulation dataset was obtained from OpenfMRI, and is publicly available; its accession number is ds000009. Note that although the full dataset includes n = 24 subjects, only 11 subjects had preprocessed data available. We therefore conducted analyses with the convenience n = 11 sample, since subject sample size was irrelevant for our purposes. Experimental design and preprocessing procedures for this dataset have been previously described in Cohen (2009).

The IAPS dataset previously used in Chang et al. (2015) was obtained from the NeuroVault whole-brain image repository (Gorgolewski et al., 2015). Images were downloaded via the NeuroVault API from the corresponding image collection (http://neurovault.org/collections/503/). The dataset contains 30 trial-level estimates for each of 172 participants (30 images in total). On each trial, participants passively viewed either a negative or a neutral IAPS image (15 of each). All methods have been previously described in Chang et al. (2015).

### Statistical modeling

#### The standard model

Consider a hypothetical fMRI experiment in which participants view 20 stimuli, half belonging to one stimulus category and half to another (as in Figure 1), and we are interested in the difference in neural response between these two categories. Let *Y*_*it*_ be the neural response of the *i*th subject at the *t*th time point in a particular voxel or region of interest (ROI), with all preprocessing already carried out and with each participant’s data separately standardized. The standard statistical model used to analyze such data is

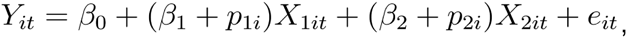

where *X*_1_ and *X*_2_ are the regressors representing the predicted neural responses toward the two categories of stimuli; *β*_0_ is the fixed intercept; *β*_1_ and *β*_2_ are the fixed effects of the two neural regressors; *p*_1_ and *p*_2_ are normally distributed participant effects of the neural regressors, representing stable subject-to-subject variability in the degree of neural response toward both stimulus types; and the *e* terms are normally distributed, observation-level errors with an AR(2) covariance structure. The regressors *X*_1_ and *X*_2_ are formed by summing over the predicted neural responses toward the individual stimuli comprising each stimulus category,

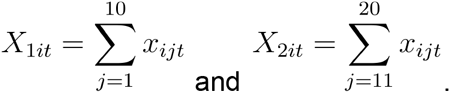

where *x*_*ijt*_ is the predicted neural response of the *i*th participant for the *j*th stimulus at time *t*. These predicted neural responses are based on convolving a stimulus presentation sequence with a hemodynamic response function (Poldrack, Mumford, & Nichols, 2011).

#### Random Stimulus Model (RSM)

The standard model posits that the stimulus-level regressors are all identical in amplitude, differing only in their presentation times (see Figure 1A)—a dubious assumption in most fMRI studies. In the RSM, we relax this assumption and allow the stimulus-level regressors to have distributions of amplitudes that are to be estimated from the data (see Figure 1B); these amplitudes are common over subjects, but vary randomly per stimuli. To achieve this, we add a set of terms 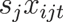 to the model, where the 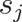 are normally distributed stimulus effects, representing stable stimulus-to-stimulus variability in the strength of the neural response. The resulting model is

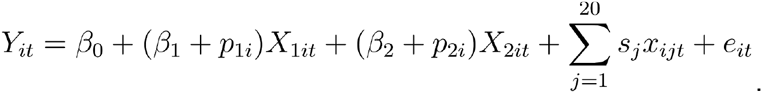

This model cannot be fit using standard mixed modeling statistical packages, such as Ime4 in R or SAS PROC MIXED, because these packages assume that each row of the dataset is associated with one and only one level of each random factor (i.e., a single participant and a single stimulus)^1^. However, for the RSM, the measurements at each time point are influenced not by a single random stimulus effect 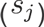, but rather by *all* of the random stimulus effects (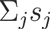). Despite this complication, it is relatively straightforward to fit the RSM as a Bayesian model using a probabilistic programming framework such as BUGS, JAGS, or Stan. For the models in this paper, we used the PyMC3 Python package (Patil, Huard, & Fonnesbeck, 2010; Salvatier, Wiecki, & Fonnesbeck, 2015), which is built on the Theano deep learning package (Bastien et al., 2012; Bergstra et al., 2010) and implements the state-of-the-art No U-Turn MCMC Sampler (Hoffman & Gelman, 2014). An alpha version of our *NiPyMC* analysis package is available online (github.com/PsychoinformaticsLab/nipymc). In Appendix 1 we give the full statistical details of the specific models we estimated in our reanalyses, including the precise distributional assumptions and the variations of the basic model that we applied to each individual dataset.

We note that there are various specification options that could be applied to the standard model and RSM described here and in Appendix 1--for example, different choices of HRF, autocorrelation parameters, motion correction, image realignment, and so on. While such options can certainly impact overall data quality and test statistics (cf. Carp, 2012) they are extremely unlikely to affect the central conclusions supported by the present results. To exert a non-negligible impact on our results, these specification options would need to have very different impacts on the standard model and RSM (otherwise the extensions would simply lead to the test statistics from both models increasing or decreasing more or less in unison, leaving their relative differences essentially unchanged). We are aware of no a priori reasons to expect this to be the case for any of the methodological procedures employed with any frequency in the literature, and reiterate that comparably large decreases in test statistics have been repeatedly observed in other domains of psychology when including random stimulus effects (Judd et al., 2012; Wolsiefer, Westfall, & Judd, 2016).

### Simulations

We conducted an extensive series of simulations in order to validate and to better understand the properties of our proposed RSM. Our first goal was to verify that the RSM could adequately recover true parameter values. Our second goal was to identify the conditions under which using a RSM produces the greatest attenuation of test statistics compared to the standard model. In orthogonal, ANOVA-like designs where the appropriate RSM can be fit in standard mixed modeling software, it can be shown that the test statistic for the standard model that ignores stimulus variability will be inflated by a factor of roughly 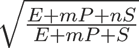, Where *n* is the number of participants, *m* is the number of stimuli, and *E*, *P*, and *S* are, respectively, the error variance, participant variance, and stimulus variance (the exact expression depends on the experimental design). While we cannot safely assume that the more complicated fMRI RSM will follow a similar inflation factor, this does give us several hypotheses about the qualitative conditions under which we should expect the worst inflation in fMRI data. Specifically, the degree of inflation should increase with participant sample size, decrease with stimulus sample size, and increase with stimulus variability. In Appendix 1 we describe the results of our simulations in detail. Here we summarize the basic structure of the simulations and their results.

In each run of the simulation, we generated data according to the RSM for a block-design experiment involving participants responding to stimuli nested in two stimulus categories. The test of interest in these simulated experiments is the difference in the fixed regression coefficients for the two stimulus categories (i.e., whether there is greater activation for one stimulus category than for the other). We varied three primary factors in our simulations: the participant sample size (*n* = 16, 32, or 64), the stimulus sample size (*m* = 16, 32, or 64), and the degree of random stimulus variability (zero, moderate, or high). Note that when the random stimulus effects have zero variance, the RSM is statistically equivalent to the standard model. We included this condition in order to investigate the performance of the RSM when the standard model is the correct model. For each simulated experiment, we fit four statistical models: the standard model, the RSM, the standard SPM-style “summary statistics” model, and a fourth model that we call the Fixed Stimulus Model, which we describe in Appendix 1. Here we focus on comparing the performance of the standard model and RSM (though, in practice, the three non-RSM models all display essentially indistinguishable behavior across all simulations).

When the true data generating process contained zero stimulus variability, the standard model and RSM yielded very similar test statistics for the stimulus category difference, for all participant and stimulus sample sizes. The exception was that when the stimulus sample size was small (*m* = 16), the test statistics from the RSM were slightly attenuated (by about 18%) compared to the standard model test statistics. However, this attenuation disappeared with increasing stimulus sample size, so that at *m* = 32 and *m* = 64, the test statistics from the two models were essentially identical, as should be the case given the lack of true stimulus variability. This provides evidence that the reduced test statistics observed in our reanalyses of real datasets (most of which were based on a sample of 100 participants) are not simply the result of the RSM always yielding lower test statistics. Instead, the RSM tends to yield lower test statistics when they should in fact be lower, namely, when there is random stimulus variability in the data that is ignored by the standard model.

When the true data generating process contained moderate stimulus variability, the RSM yielded consistently lower test statistics than the standard model. This reduction was exacerbated when the stimulus sample size was small (*m* = 16) and attenuated somewhat when the stimulus sample size was large (*m* = 64). The opposite pattern held for the participant sample size: the reduction in the RSM test statistic was largest when the participant sample size was large (*n* = 64) and smallest when the participant sample size was small (*n* = 16). These patterns are consistent with what has been observed in previous behavioral work (Judd et al., 2012). In the best case (*n* = 16 and *m* = 64) the RSM test statistics were still reduced by an average of 17%compared to the standard model test statistics. In the worst case (*n* = 64 and *m* = 16), the RSM test statistics were reduced by an average of 67%. Finally, when the true data generating process contained high stimulus variability, the same qualitative patterns held, but the reduction in test statistics was even greater, ranging from 41% in the best case to 81% in the worst case.

### Literature review

For our survey of the literature using task-based fMRI, we randomly selected published papers listed in the Neurosynth database -- skipping over studies that used resting-state fMRI --and coded each study on a small set of criteria until we had reached exactly 100 experiments. Four of the papers we sampled described two experiments, one paper described three experiments, and the rest described a single experiment, so that the 100 experiments we sampled came from 94 unique papers. We coded each experiment for (a) whether the stimuli were crossed with participants or nested in participants, (b) the type of stimuli used, (c) the total number of stimuli, (d) the number of stimulus categories, and (e) whether the study was eligible to have applied a RSM to the data obtained from the study. Generally speaking, experiments were deemed eligible to have applied a RSM if (i) the experiment used more stimuli than stimulus categories (so that the individual stimulus effects are statistically identifiable), and (ii) the sampled stimuli could not be considered to fully exhaust the population of stimuli over which generalization was intended. In the handful of cases where it was not totally clear from the text whether a RSM could have been applied, we decided to err on the conservative side and deem the study ineligible. A spreadsheet with the detailed study-level results of our survey can be found at github.com/PsychoinformaticsLab/nipymc. We ultimately found that 63/100 of the experiments (95% Jeffreys interval = [53%, 72%]) were eligible to have applied a RSM, and thus the published test statistics for these experiments are likely inflated relative to the more appropriate RSM test statistics.

## Acknowledgements

This work was supported by National Institutes of Health Grant R01MH096906.

## Appendix 1

### Statistical models

As mentioned in the main text, we implemented the Random Stimulus Model (RSM) as a Bayesian model, primarily in order to circumvent certain limitations in standard statistical software for fitting linear mixed models. In order to facilitate comparisons between the RSM and the standard model applied in most research papers, we also implemented the standard model as a Bayesian model, so that the two models differed only in their inclusion of random stimulus effects. Here we give the full Bayesian mixed models that we used for all 6 datasets we reanalyzed. Each model contains a set of stimulus-level regressors 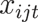 that, as in the main text, represent the predicted neural response of the *i*th subject toward the *j*th stimulus at the *t*th time point. These regressors were constructed by convolving a stimulus presentation sequence, indicating when a stimulus was being presented, with a double-gamma hemodynamic response function (HRF) with parameters fixed at the default values used in the SPM package.

For each task, the model was applied separately to 100 regions-of-interest (ROIs) defined via meta-analytic k-means clustering of over 11,000 studies in the Neurosynth dataset (http://neurosynth.org; details are reported in a forthcoming publication). The 100-ROI whole-brain cluster map is available from the project GitHub repository (http://github.com/PsychoinformaticsLab/nipymc). Note that this mask was used purely for convenience, and the results reported here should not depend in any way on the the choice of ROIs (as can be seen from the near ubiquitous drop in test statistics across all ROIs in Fig. 4).

#### HCP Emotion model

The full Bayesian mixed model for *Y*_*it*_, representing the neural response within a particular ROI (separately standardized for each subject) for the *i*th subject toward the *j*th stimulus in the *k*th experimental condition at the *t*th time point, is

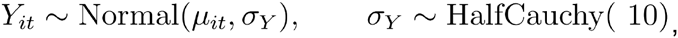

where HalfCauchy was chosen as the default, weakly informative prior for the standard deviation (Gelman, 2006), and the mean is

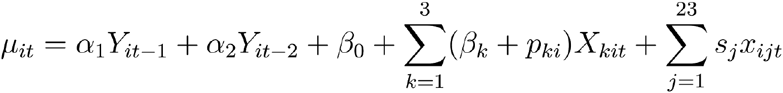

Where *α*_1_ and *α*_2_ are fixed effects accounting for autoregressive order-2 variation; *β*_0_ and *β*_*k*_ are fixed effects of the intercept and condition *k,* respectively; *p*_*ki*_ are the random subject effects, the disturbance from *β*_*k*_ for subject *i,* condition *k*; ^*s*_*j*_^ are the random stimulus effect for stimulus *j*. The autoregressive fixed effects have Cauchy priors and the remaining fixed effects have Normal priors,

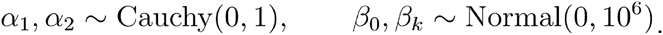

The random subject effects are Normal with the same HalfCauchy prior

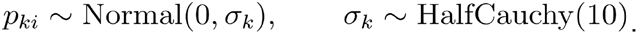

The stimuli random effects likewise have the same type Normal/HalfCauchy priors; for shape stimuli,

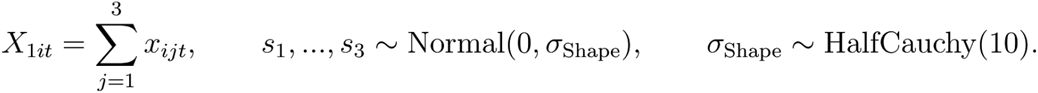

For anger face stimuli,

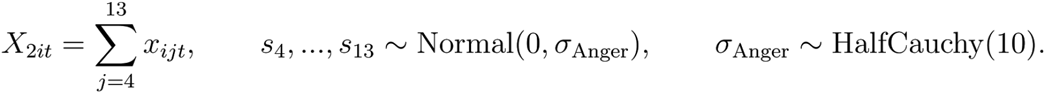

For fear face stimuli,

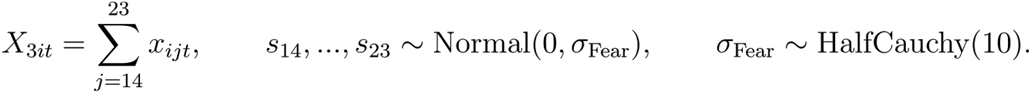

The two contrasts of interest in this model are the Face vs. Shape contrast, 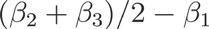, and the Anger vs. Fear contrast, *β*_2_ – *β*_3_.

#### HCP Working Memory model

The full Bayesian mixed model for *Y*_*it*_ -- representing the neural response within a particular ROI (separately standardized for each subject) for the *i* th subject toward the *j*th stimulus in the *k*th stimulus category under the *l*th N-back condition at the tth time point -- has a similar form as above:

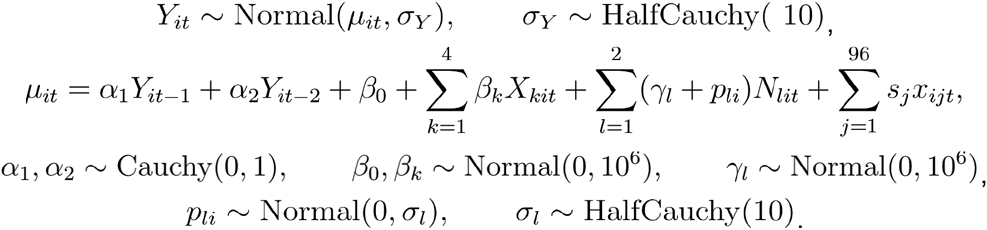

For this model, the stimuli are as follows; for body stimuli,

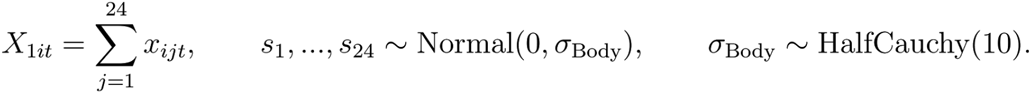

For place stimuli,

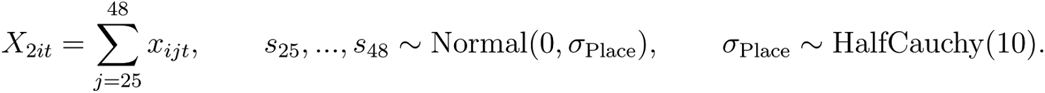

For face stimuli,

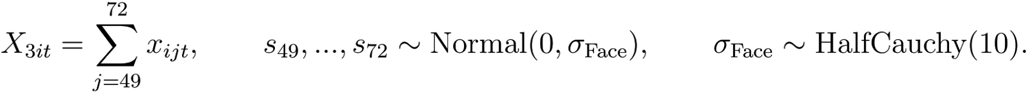

For tool stimuli,

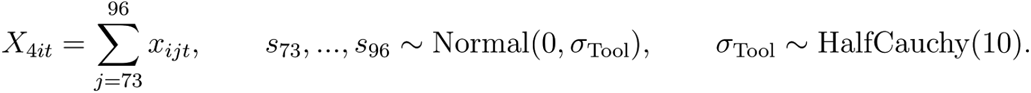

The regressors *X*_l*it*_ to *X*_4*it*_ were constructed in the usual way (by convolution with a double-gamma HRF) and then orthogonalized with respect to *N*_l*it*_ and *N*_2*it*_, which were the regressors of interest. Each in the series of 24 trials consists of both 1- and 2-back stimuli. *N*_1*it*_ and *N*_2*it*_ represent the predicted neural responses for the 0-back and 2-back trials, respectively, of the N-back task used in the HCP Working Memory data. Note that both the participants and the stimuli were crossed with the two N-back conditions, i.e., all subjects made responses in both conditions and all stimuli received responses under both the conditions. The contrast of interest in this model is the difference in response between the 0-back and 2-back conditions, *γ*_1_ – *γ*_2_.

#### HCP Social Cognition model

The full Bayesian mixed model for *Y*_*it*_, representing the neural response within a particular ROI (separately standardized for each subject) for the *i*th subject toward the *j*th stimulus in the *k*th experimental condition at the *t*th time point, has the form:

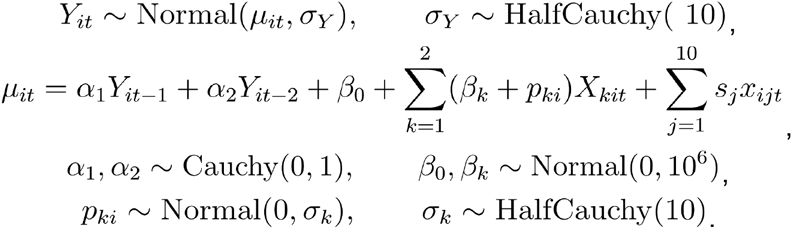

The stimuli are as follows; for the random stimuli,

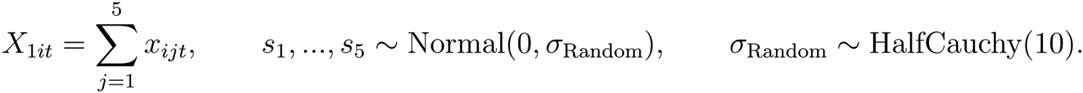

For the mental stimuli:

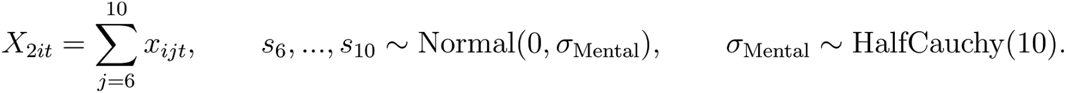

The contrast of interest in this model is the random vs. mental contrast, *β*_1_ – *β*_2_.

#### Chang et al. (2015) model

For this dataset we used a slightly simplified, approximate version of the full RSM, quite similar to what Gelman & Hill (2007) refer to as a “no-pooling model.” The basic idea is that rather than directly modeling all of the neural data for all subjects for a particular ROI at once, one first fits separate subject-level models that simply contain a separate fixed regressor for each stimulus. Since the participants and stimuli are fully crossed in this dataset, this produces *n × m* estimated regression coefficients, where *n* is the number of subjects and *m* is the number of stimuli. These estimated regression coefficients are then propagated upward to serve as the data in a higher-level RSM that includes crossed random participant and stimulus effects. This model is only an approximate version of the full RSM, as it ignores the varying degrees of uncertainty in the stimulus-level coefficients estimated at the first level (due in particular to factors such as differences in how often the stimuli were presented and how collinear the stimulus-level regressors were for each subject). We used this approximate method because this is the form in which the data were made available on the public NeuroVault repository (NeuroVault.org; Gorgolewski et al., 2015). However, it is worth noting that two practical advantages of this simplified model are that (i) it can be estimated relatively quickly and efficiently compared to the full RSM, and (ii) it can be fit using standard mixed modeling software such as R’s Ime4 package or SAS PROC MIXED, since the higher-level model is just a straightforward crossed random effects model with one and only one stimulus effect associated with each data point.

For the statistical details of the first-level model, which was applied separately to each subject’s neural data, see Chang et al. (2015). From these first-level models one obtains *n × m* coefficients *b*_*ij*_ representing the estimated regression coefficient for the neural response of the *i*th subject toward the *j*th stimulus. After standardizing the coefficients separately for each subject, the 2nd level model is then, as before, the Normal/HalfCauchy observation model

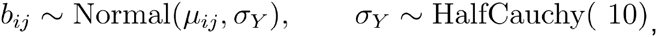

with mean given by

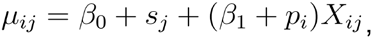

where now *β*_0_ is the mean response to all stimuli, *S*_*j*_ is the random stimulus effect, *β*_1_ is the fixed effect of valence, *p*_*i*_ is the random subject effect of valence, and *X*_*ij*_ is a dummy variable indicating whether the *j*th stimulus photograph was negative or neutral in valence. The fixed variables have wide Normal priors,

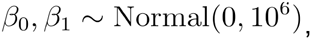

And the random effects have the Normal/HalfCauchy priors as before:

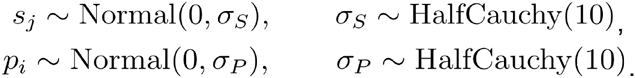

#### OpenfMRI Emotion Regulation model

The full Bayesian mixed model for *Y*_*it*_, representing the neural response within a particular ROI (separately standardized for each subject) for the *i*th subject toward the *j*th stimulus in the *k*th experimental condition at the *t*th time point, takes the form of the time-series models above:

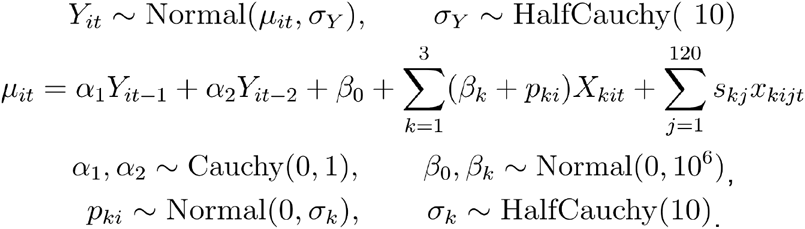

For the Negative/Suppress stimuli,

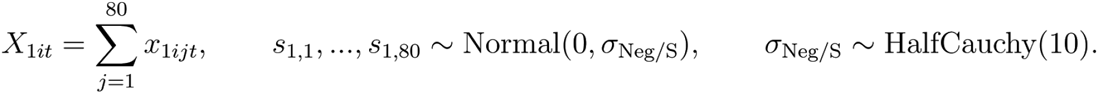

For the Negative/Attend stimuli,

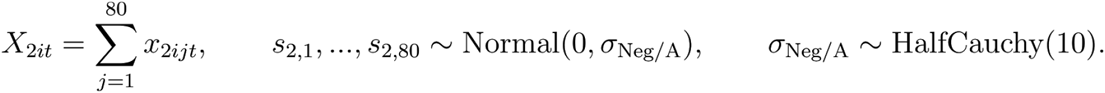

For the Neutral/Attend stimuli,

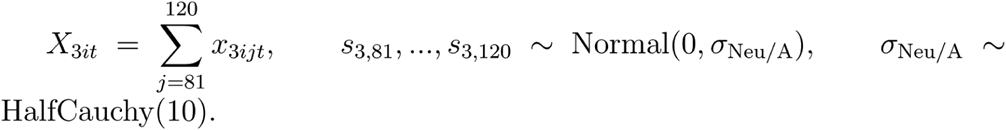

Note that all of the negative stimuli (*j* = 1,…, 80) are observed in both the Suppress and Attend conditions, while the neutral stimuli (*j* = 81,…, 120) are observed only in the Attend condition, so that there are a total of 200 stimulus effects (80+80+40) for the 120 stimuli. The two contrasts of interest in this model are the Negative vs. Neutral contrast, 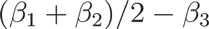 and the Suppress vs. Attend contrast, 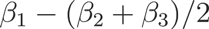. Although we report only the Negative vs. Neutral results in the main text for the sake of brevity, the results for the Suppress vs. Attend contrast were nearly identical in terms of the impact of the RSM on test statistics; test statistics in the RSM were reduced by about 14% on average compared to those from the standard model.

#### HCP Language model

This dataset consisted of subjects’ responses toward 6 stories and 831 math problems (drawn from a much larger potential set of over 7,000 unique stimuli). These math problems were generally presented a very small number of times each--most appeared only once in the dataset, and many appeared only twice. Due to the intensive computation that would have been required to estimating models with 831 extra parameters (most of which were either not identifiable or only weakly identifiable), we decided to group all of the math stimuli with fewer than 3 responses into a single “dummy” stimulus, resulting in a new total of 272 math stimuli. The full Bayesian mixed model for *Y*_*it*_, representing the neural response within a particular ROI (separately standardized for each subject) for the *i*th subject toward the *j*th stimulus in the *k*th experimental condition at the *t*th time point, takes the same form as the previous model:

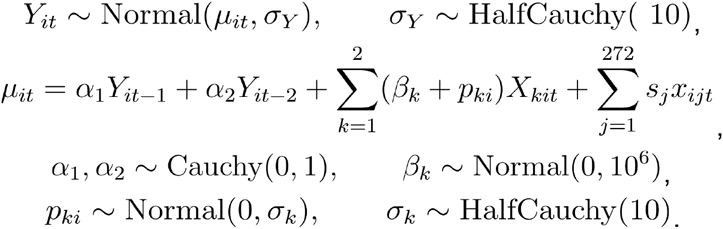

For the story stimuli,

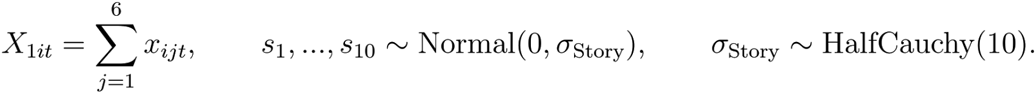

For the math stimuli,

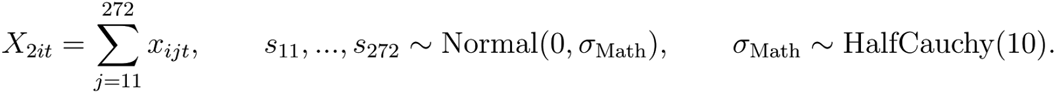

The contrast of interest in this model is the Story vs. Math contrast, *β*_1_−*β*_2_

### Simulation

We simulated experiments in which *n* participants respond one time each to *m* stimuli that belong to one of two stimulus categories, and the contrast of interest is the difference in neural activation between the two categories. The simulated experiments used a block design with the stimuli presented in alternating blocks of 8, with the order of the stimulus category presentations (i.e., whether the sequence began with category A or category B) counterbalanced across participants. Each experimental run consisted of an initial stimulus presented for 1 second, then a 2-second inter-stimulus interval (ISI), then another stimulus presented for 1 second, then another 2-second ISI, and so on until all stimuli had been presented once. The stimulus regressors 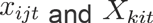 were formed in the same way as described in the previous “Statistical models” section.

The data *Y*_*it*_, representing the neural response within a particular voxel or ROI for the *i*th subject toward the *j*th stimulus in the *k*th experimental condition at the *t*th time point, were generated according to the model

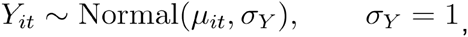

where the mean is given by

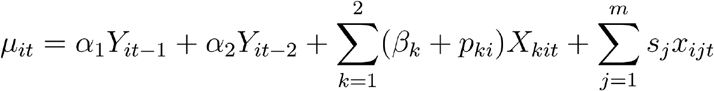

The autocorrelation parameters were set equal to

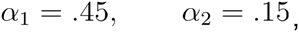

and the fixed means for the two stimulus categories were set equal to

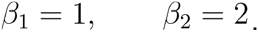

The random participant effects *p*_*ki*_ and the random stimulus effects 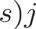 were distributed as

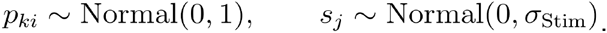

In our simulations we varied the number of participants *n* ∈{16, 32, 64}; the number of stimuli *m* ∈ {16, 32, 64}; and the standard deviation of the random stimulus effects *σ*_Stim_ ∈ {0, 1, 2}. We ran 500 iterations of the simulation within each of these parameter combinations. In each iteration of the simulation we fit 4 statistical models to the simulated dataset, which we describe next.

#### SPM model

The first model that we fit to each simulated dataset is perhaps the most widely used statistical model in neuroimaging, serving as the default model in the SPM software package. In practice it is implemented in a two-stage procedure. In the first stage, one fits separate regression models to the data from each participant, where the model for the *i*th participant is

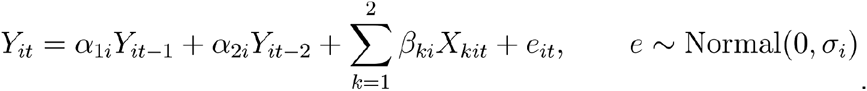

In the second stage, one computes the difference 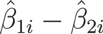 for each participant and then performs a one-sample t-test on these differences. This model can be seen as an approximation to a mixed-effects model with random participant effects -- what we refer to below as the No Stimulus Model (NSM) and in the main text as the standard model -- valid to the extent that the intrasubject variances are all equal. We include the results of this model in our simulations simply to show that it yields practically equivalent results to the NSM model that we focus on in the main text, despite their slight differences in assumptions. In the main text we focus on comparing the Random Stimulus Model (RSM) to the NSM, rather than the SPM model, because it is a cleaner comparison, since the RSM and NSM are identical except for their inclusion of random stimulus effects.

#### Fixed stimulus model (FSM)

The second model that we fit is similar in structure to the RSM, except the stimuli are treated as fixed effects rather than random effects. In other words, a separate fixed regression coefficient is estimated for each individual stimulus regressor. This model has been described by (Mumford, Turner, Ashby, & Poldrack, 2012), who referred to it as the “Least Squares -- All”model. Specifically, the FSM that we estimated in each iteration is

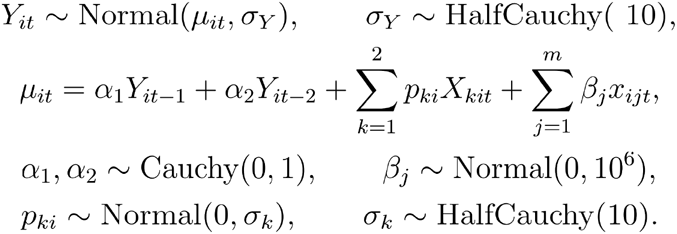

Note that the random stimulus effects *s*_*j*_ are replaced with fixed effects *β*_*j*_, which also requires that the category fixed effects *β*_1_ and *β*_2_ are removed from the model, since they would be perfectly collinear with the fixed stimulus effects. The contrast between the two stimulus categories is computed as the mean of the *β*_*j*_ in stimulus category B minus the mean of the *β*_*j*_ in stimulus category A.

#### No stimulus model (NSM)

This is the model that we referred to in the main text as the “standard model.” Unlike the FSM and RSM, the NSM does not include any stimulus effects at all, hence the name we use here. Specifically, the NSM that we estimated in each iteration is:

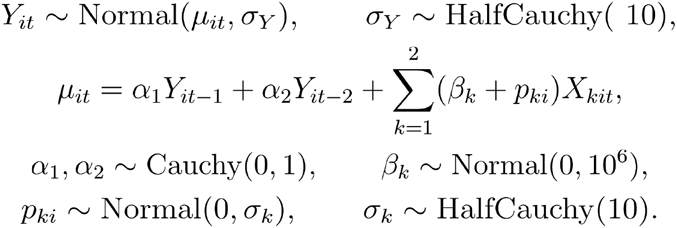

The contrast between the stimulus categories was computed as *β*_2_ − *β*_1_.

#### Random stimulus model (RSM)

The RSM that we estimated in each iteration is:

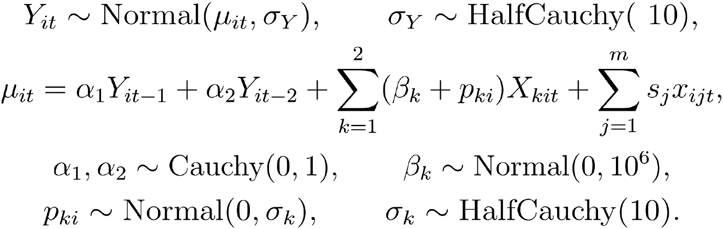

For stimulus category A,

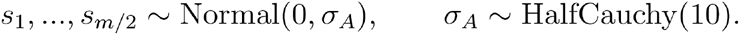

For stimulus category B,

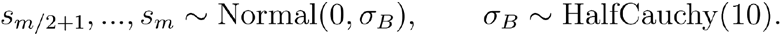

The contrast between the stimulus categories was computed as *β*_2_ – *β*_1_.

#### False positive rates under the SPM model

We first present the results for the false positive rates (i.e., Type 1 error rates) of the SPM model in a separate simulation identical to the main simulation, but with *β*_1_ = *β*_2_ = 1, so that the null hypothesis of no difference in the fixed category means is true. These error rates are based on running 500 iterations for each parameter combination, and they are computed for several levels of *α,* the decision threshold. Test levels of *α*= 0.05, 0.01, 0.005 and 0.001 are considered, which have nominal Monte Carlo 95% confidence intervals of ± 0.019, ± 0.009, ± 0.006, and ± 0.003, respectively.

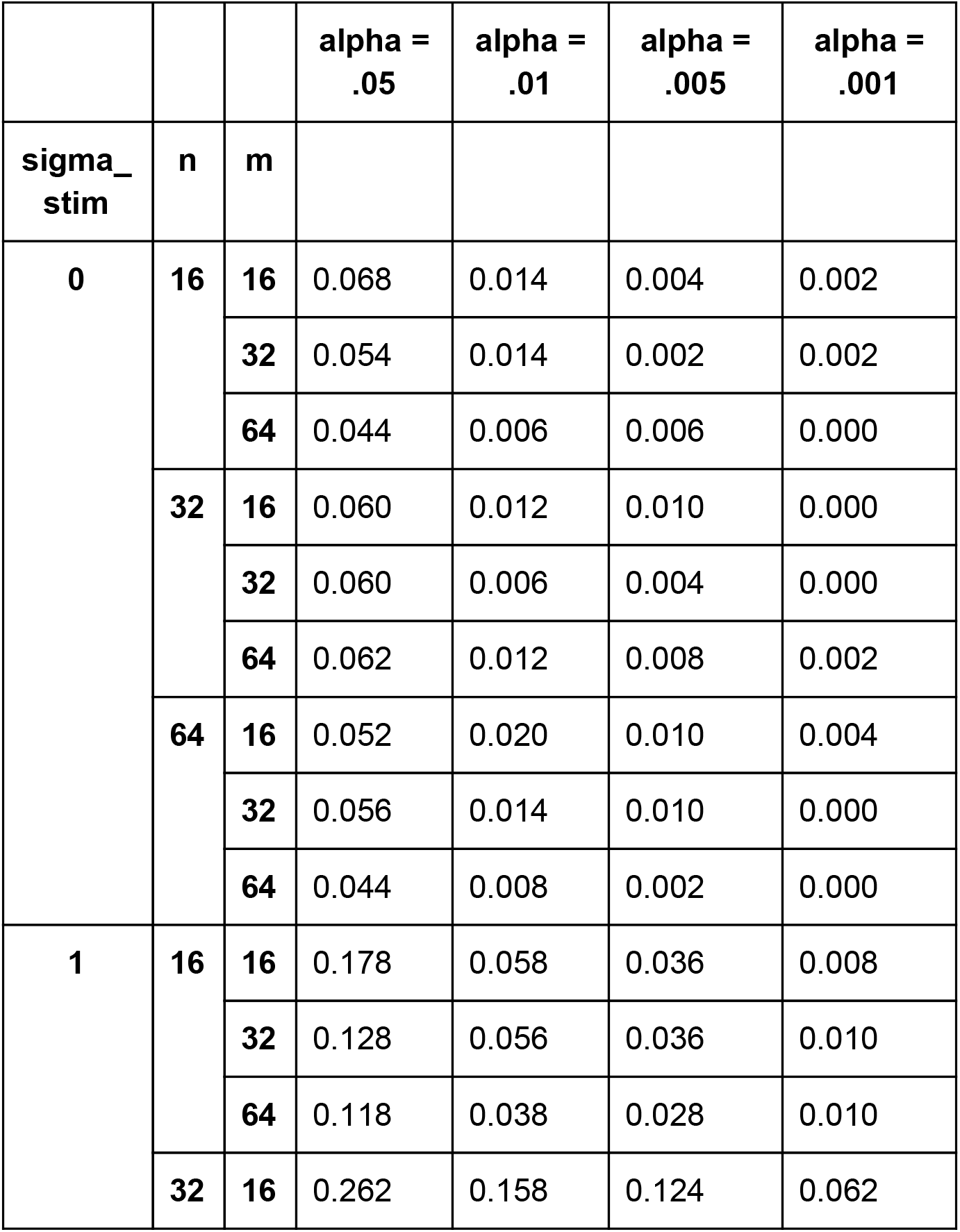

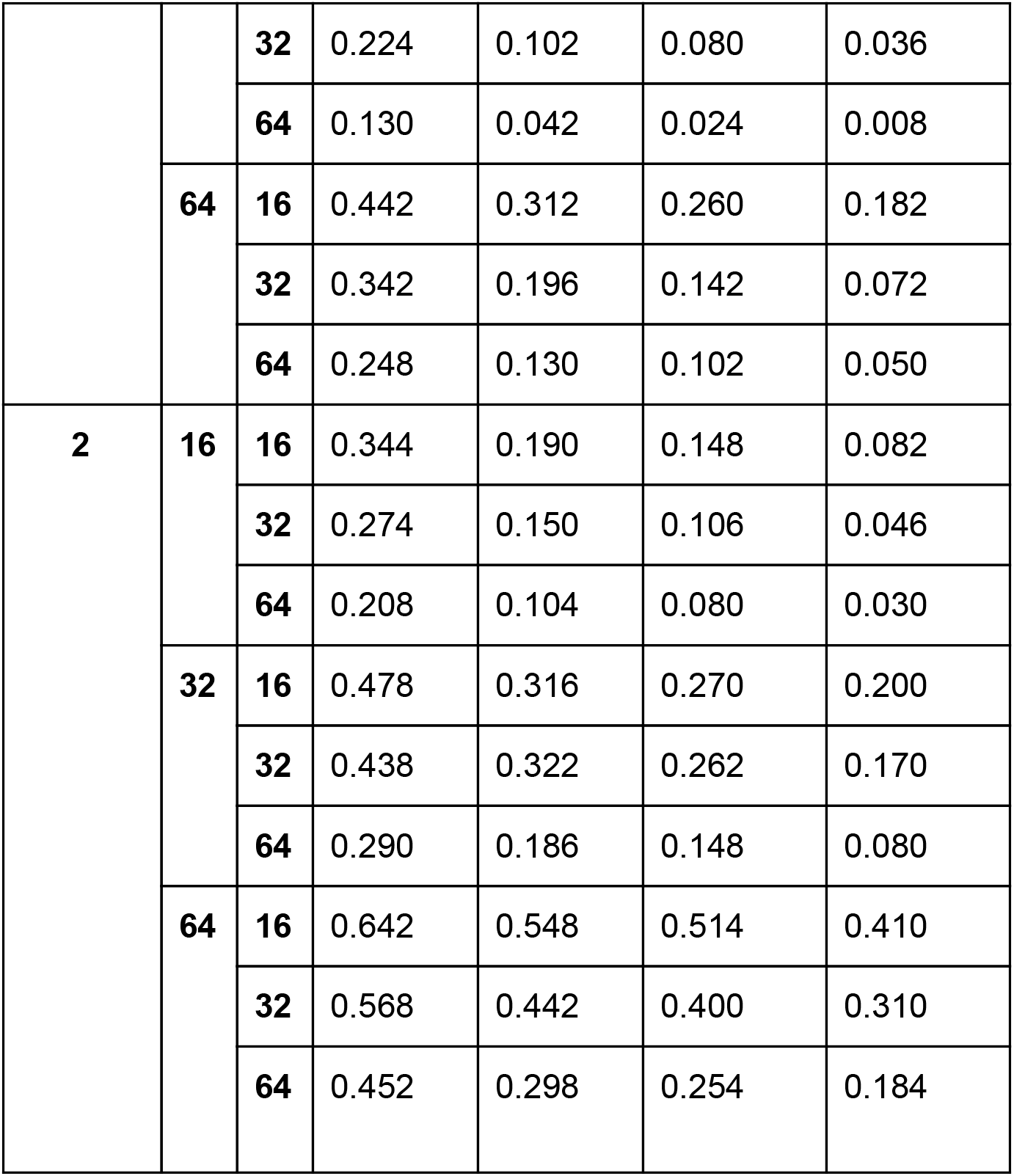

#### Inflation of test statistics

The plots below show the simulated test statistics for the contrast between the two stimulus categories for all four models in each parameter combination. The points in each panel give the mean test statistic across 500 simulation runs, and the error bars span plus-or-minus one standard deviation of the test statistics across the 500 simulation runs. It is worth noting that the parameter estimates we obtained for all three of the Bayesian models described above were not discernibly affected by the particular choice of priors. While we initially experimented with more informative priors (e.g., modeling stimulus or subject variances as ~ HalfCauchy(1)), the insensitivity of the posterior estimates to the choice of prior ultimately led us to opt for the relatively weak, or uninformative, priors detailed above. The interpretation of these simulation results is given in the Methods section in the main text. The full simulation results for all parameters of all models under all parameter combinations can be found at http://github.com/PsychoinformaticsLab/nipymc.

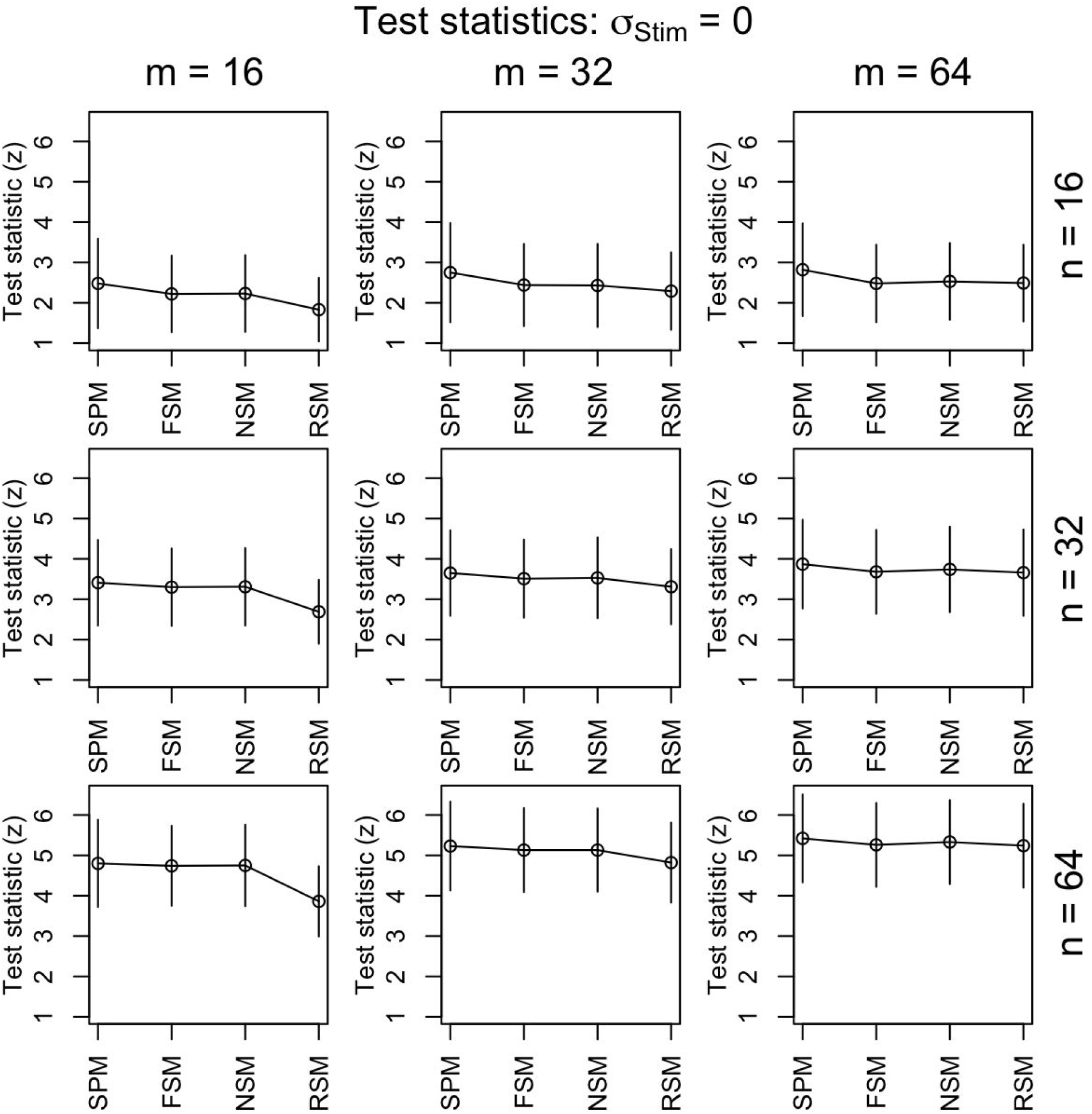

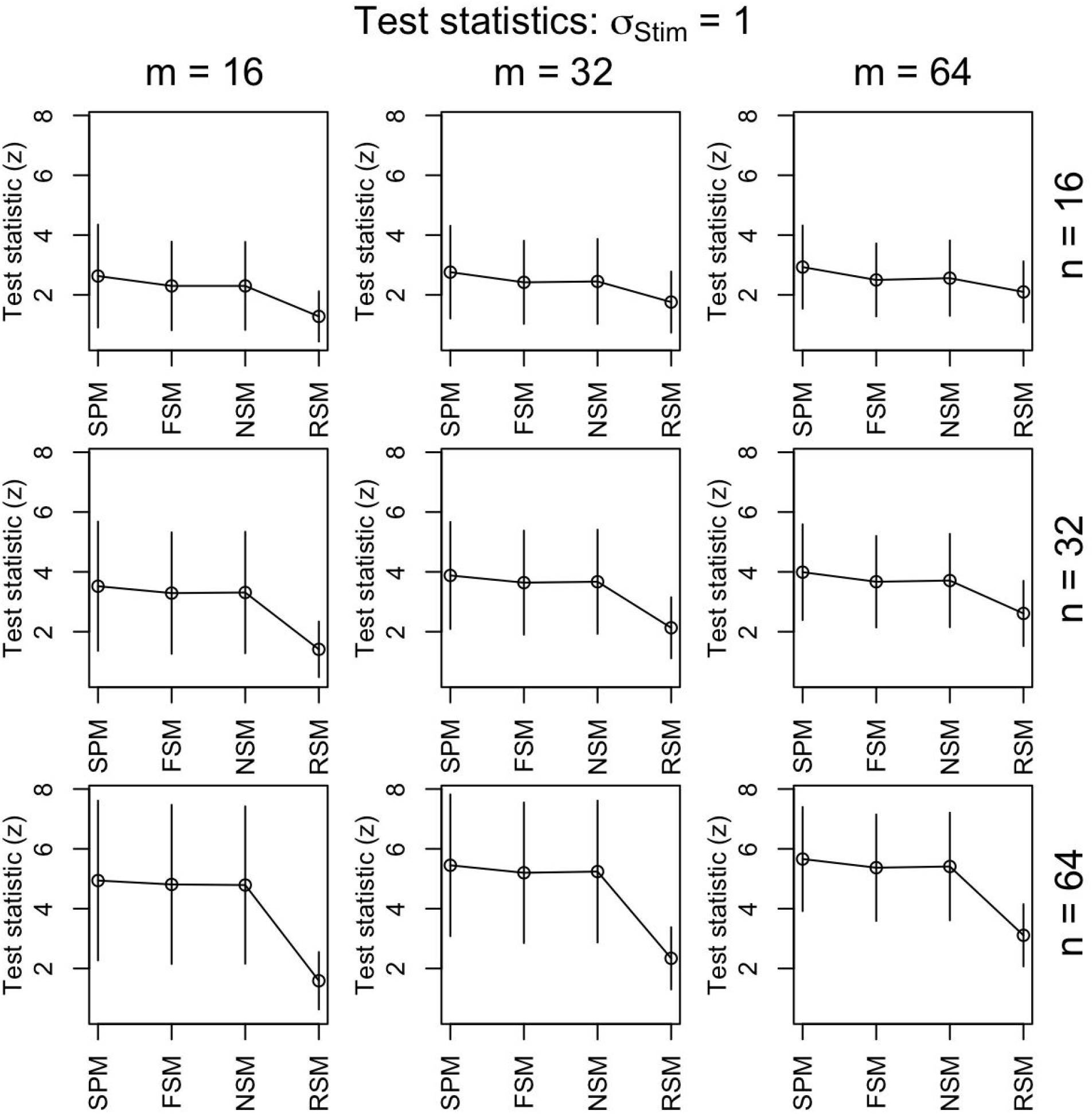

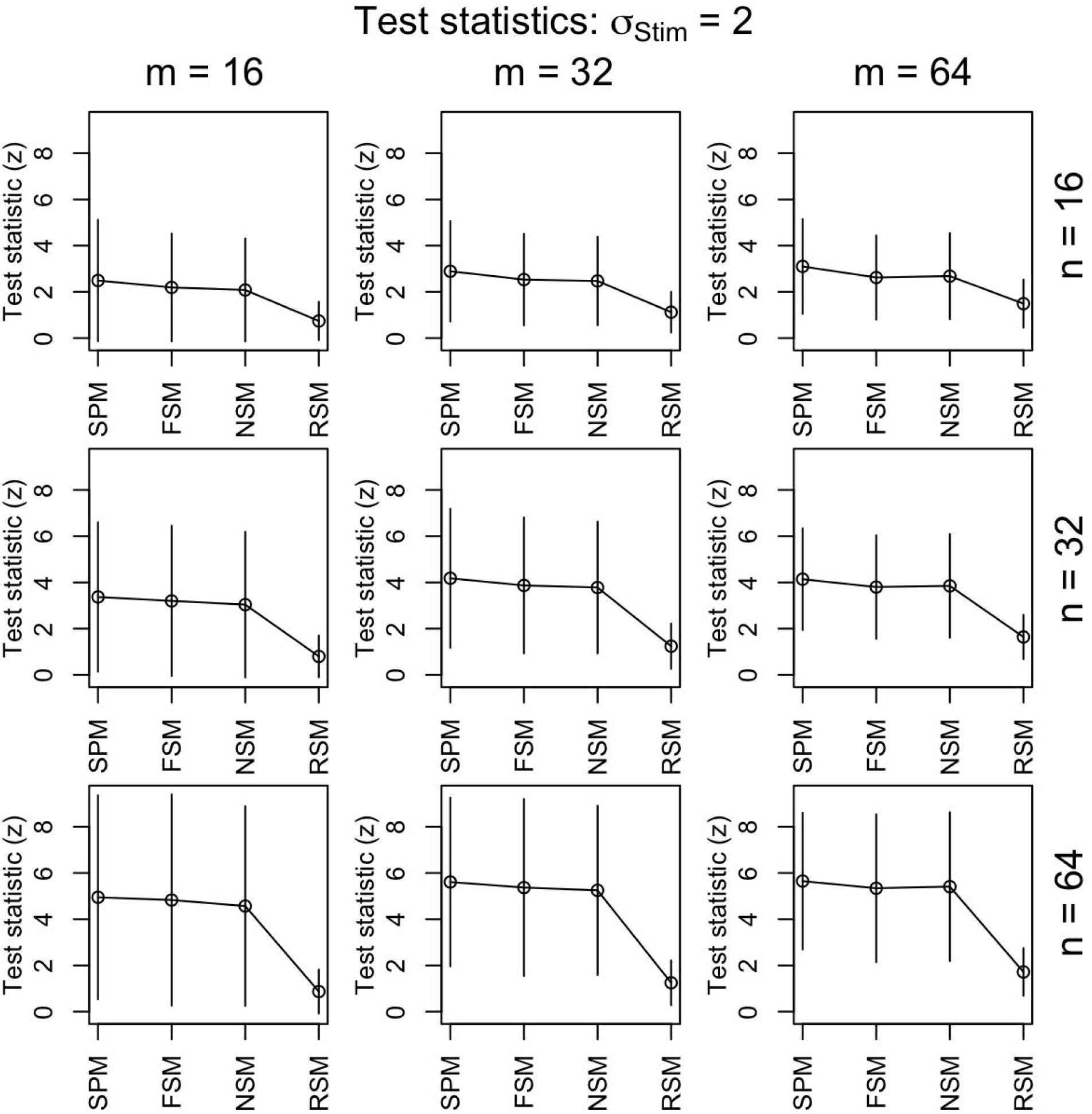

## Footnotes

1 While the full model as written here cannot be estimated in standard software due to the limitations we mentioned, it is possible to use standard software to fit a slightly simplified, approximate version of this model, quite similar to what (Gelman & Hill, 2007) refer to as a “no-pooling model.” As we discuss in Appendix 1, we did apply this approximate model to one of our reanalyzed datasets, the (Chang, Gianaros, Manuck, Krishnan, & Wager, 2015) data.

